## Supplementary figures for "Genetic mutations in GLP-1/Notch pathway reveal distinct mechanisms of Notch signaling in germline stem cell regulation"

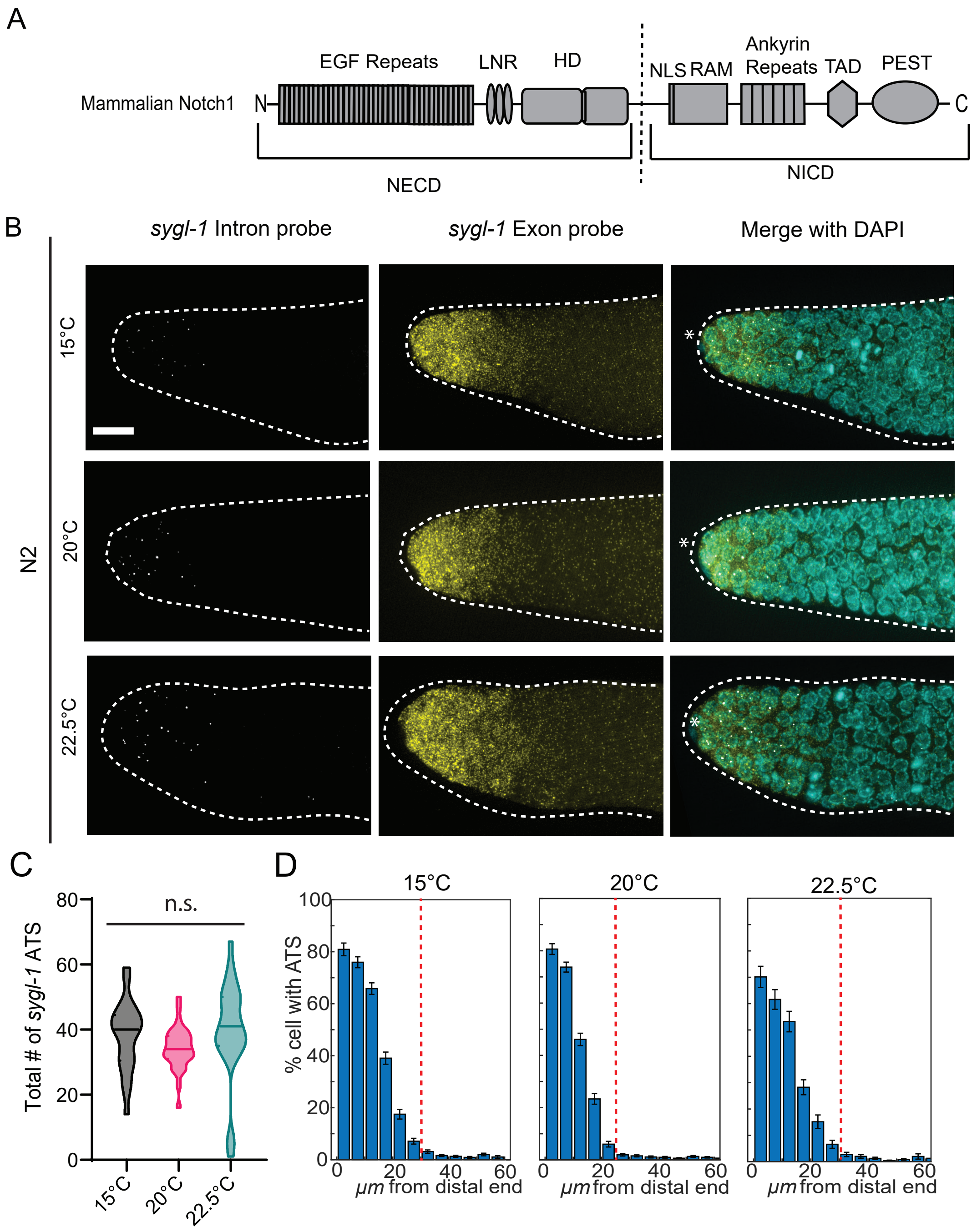
A
Mammalian Notch1
N
EGF Repeats
LNR
HD
NLS
RAM
Ankyrin Repeats
TAD
PEST
C
NECD
NICD
B
sygl-1 Intron probe
sygl-1 Exon probe
Merge with DAPI
N2
15°C
20°C
22.5°C
C
Total # of sygl-1 ATS
n.s.
80
60
40
20
0
15°C
20°C
22.5°C
D
15°C
20°C
22.5°C
% cell with ATS
100
80
60
40
20
0
0
20
40
60
μm from distal end

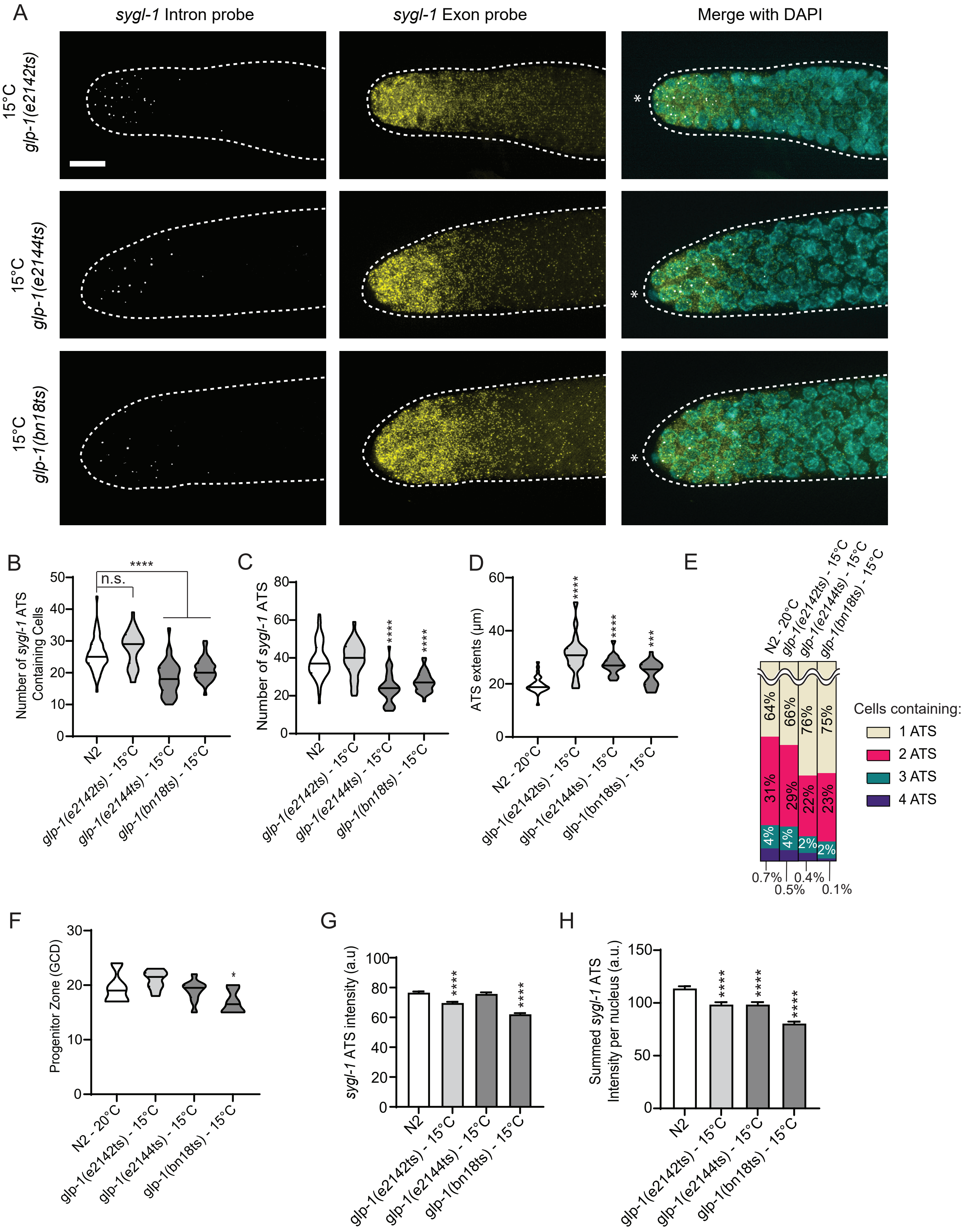
A
sygl-1 Intron probe
sygl-1 Exon probe
Merge with DAPI
15°C glp-1(e2142ts)
15°C glp-1(e2144ts)
15°C glp-1(bn18ts)
B
Number of sygl-1 ATS Containing Cells
n.s.
****
N2
glp-1(e2142ts) - 15°C
glp-1(e2144ts) - 15°C
glp-1(bn18ts) - 15°C
C
Number of sygl-1 ATS
D
ATS extents (µm)
N2 - 20°C
E
Cells containing:
1 ATS
2 ATS
3 ATS
4 ATS
64%
66%
76%
75%
31%
29%
22%
23%
4%
4%
2%
2%
0.7%
0.5%
0.4%
0.1%
F
Progenitor Zone (GCD)
G
sygl-1 ATS intensity (a.u)
H
Summed sygl-1 ATS Intensity per nucleus (a.u.)

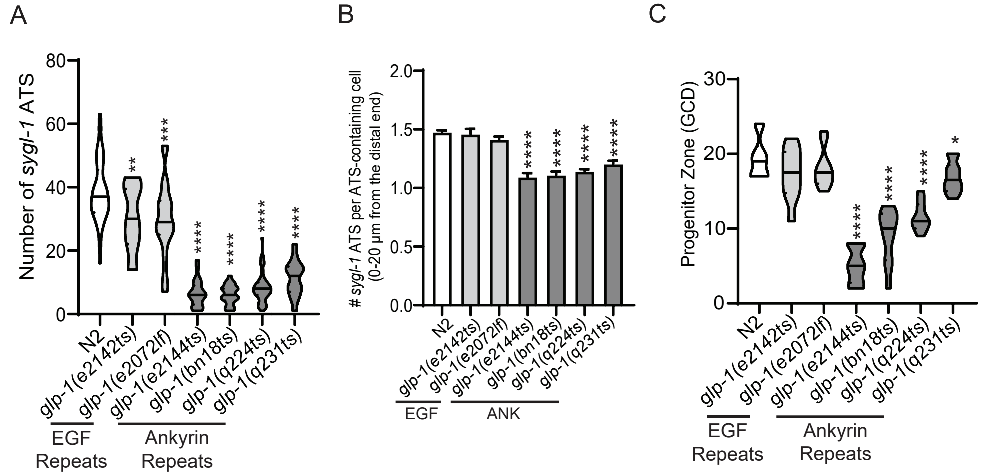
A
Number of sygl-1 ATS
80
60
40
20
0
**
***
****
****
****
****
N2
glp-1(e2142ts)
glp-1(e2072lf)
glp-1(e2144ts)
glp-1(bn18ts)
glp-1(q224ts)
glp-1(q231ts)
EGF Repeats
Ankyrin Repeats
B
### sygl-1 ATS per ATS-containing cell (0-20 μm from the distal end)
2.0
1.5
1.0
0.5
0.0
****
****
****
****
N2
glp-1(e2142ts)
glp-1(e2072lf)
glp-1(e2144ts)
glp-1(bn18ts)
glp-1(q224ts)
glp-1(q231ts)
EGF
ANK
C
Progenitor Zone (GCD)
30
20
10
0
****
****
****
*
N2
glp-1(e2142ts)
glp-1(e2072lf)
glp-1(e2144ts)
glp-1(bn18ts)
glp-1(q224ts)
glp-1(q231ts)
EGF Repeats
Ankyrin Repeats

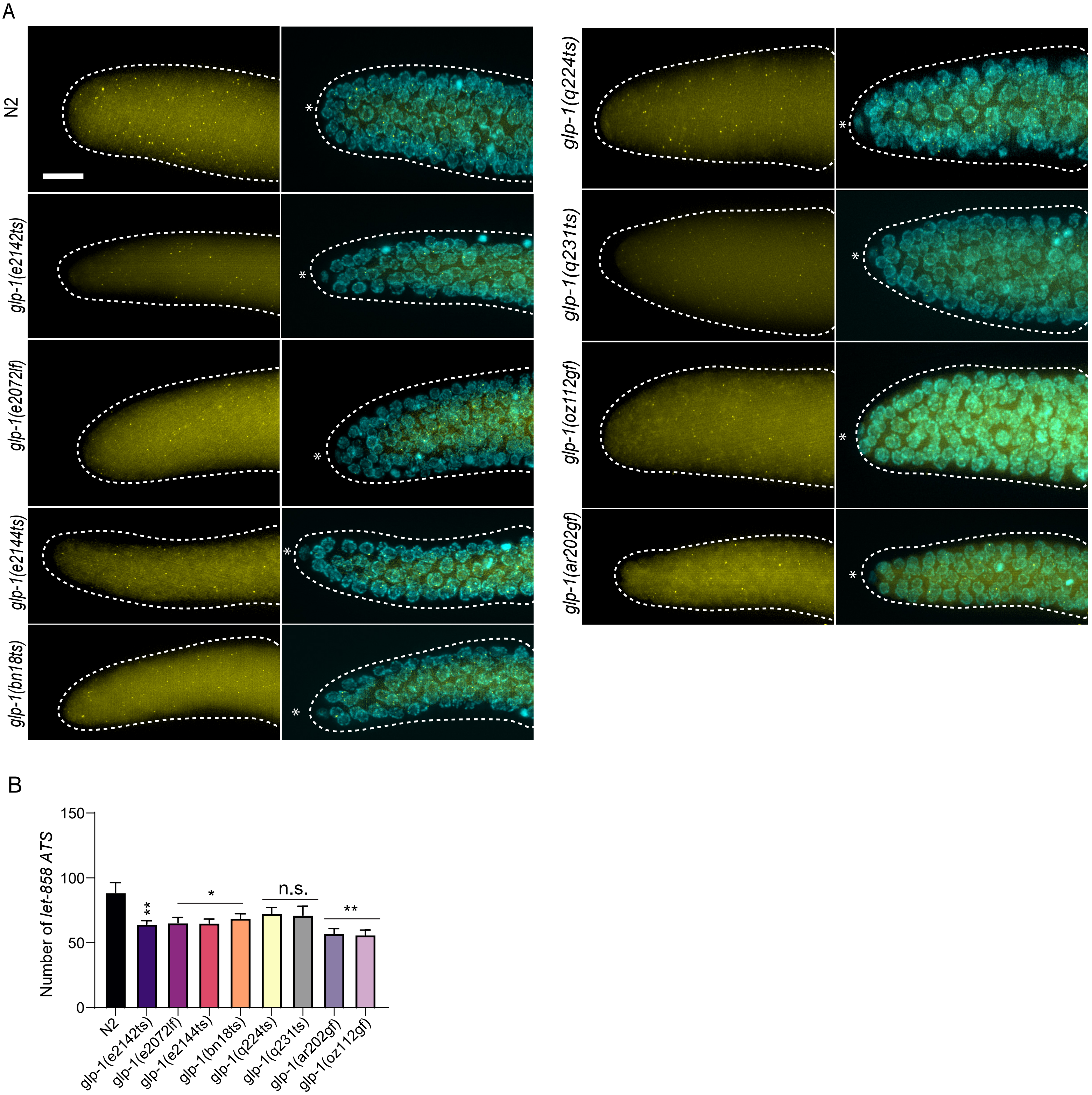
A
N2
glp-1(e2142ts)
glp-1(e2072lf)
glp-1(e2144ts)
glp-1(bn18ts)
glp-1(q224ts)
glp-1(q231ts)
glp-1(oz112gf)
glp-1(ar202gf)
B
Number of let-858 ATS
150
100
50
0
**
*
n.s.
**
N2
glp-1(e2142ts)
glp-1(e2072lf)
glp-1(e2144ts)
glp-1(bn18ts)
glp-1(q224ts)
glp-1(q231ts)
glp-1(ar202gf)
glp-1(oz112gf)

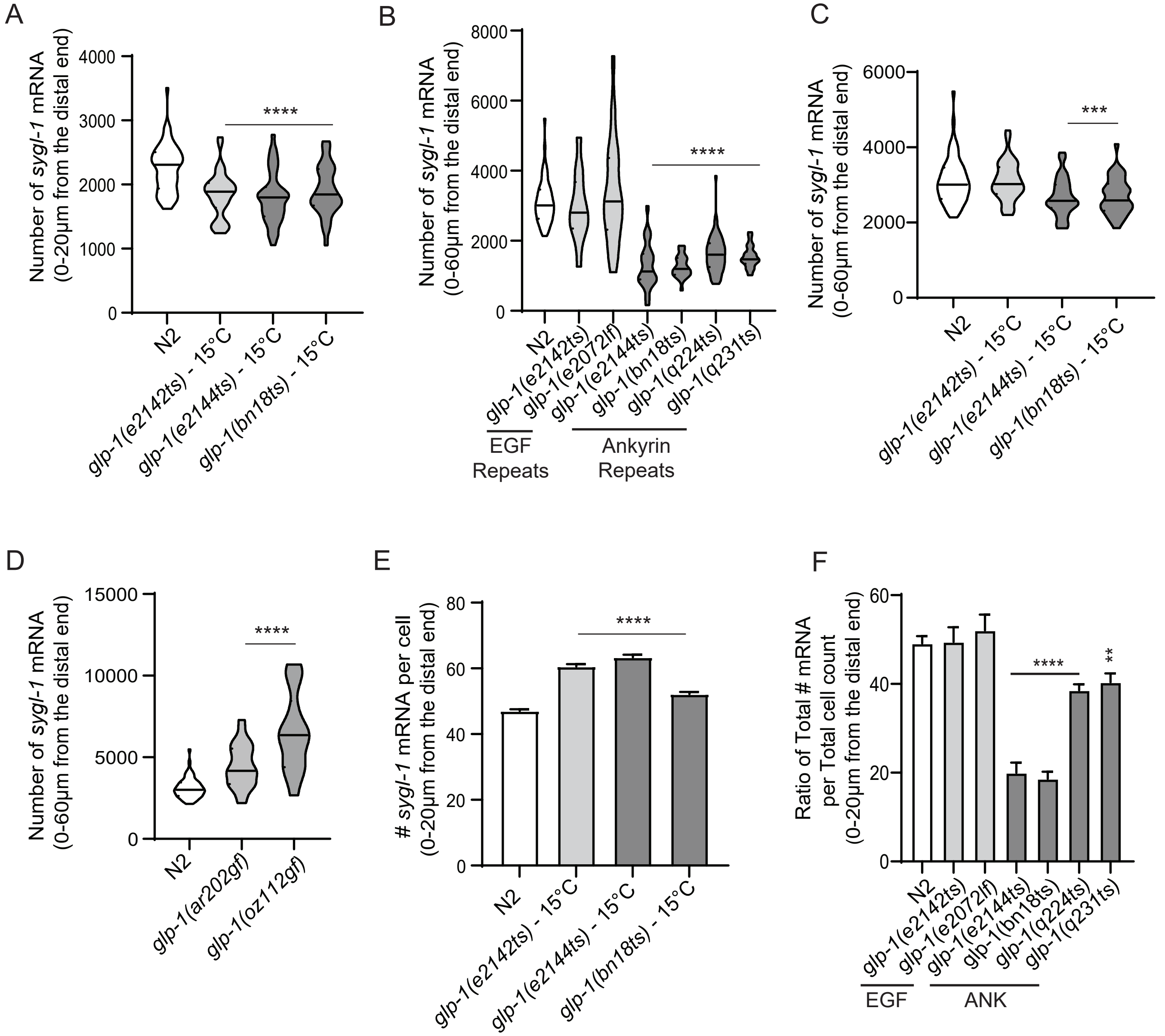

A
Number of *sygl-1* mRNA (0-20µm from the distal end)
0
1000
2000
3000
4000
****
N2
*glp-1(e2142ts)* - 15°C
*glp-1(e2144ts)* - 15°C
*glp-1(bn18ts)* - 15°C
B
Number of *sygl-1* mRNA (0-60µm from the distal end)
0
2000
4000
6000
8000
****
N2
*glp-1(e2142ts)*
*glp-1(e2072lf)*
*glp-1(e2144ts)*
*glp-1(bn18ts)*
*glp-1(q224ts)*
*glp-1(q231ts)*
EGF Repeats
Ankyrin Repeats
C
Number of *sygl-1* mRNA (0-60µm from the distal end)
0
2000
4000
6000
***
N2
*glp-1(e2142ts)* - 15°C
*glp-1(e2144ts)* - 15°C
*glp-1(bn18ts)* - 15°C
D
Number of *sygl-1* mRNA (0-60µm from the distal end)
0
5000
10000
15000
****
N2
*glp-1(ar202gf)*
*glp-1(oz112gf)*
E
### *sygl-1* mRNA per cell (0-20µm from the distal end)
0
20
40
60
80
****
N2
*glp-1(e2142ts)* - 15°C
*glp-1(e2144ts)* - 15°C
*glp-1(bn18ts)* - 15°C
F
Ratio of Total # mRNA per Total cell count (0-20µm from the distal end)
0
20
40
60
****
**
N2
*glp-1(e2142ts)*
*glp-1(e2072lf)*
*glp-1(e2144ts)*
*glp-1(bn18ts)*
*glp-1(q224ts)*
*glp-1(q231ts)*
EGF
ANK

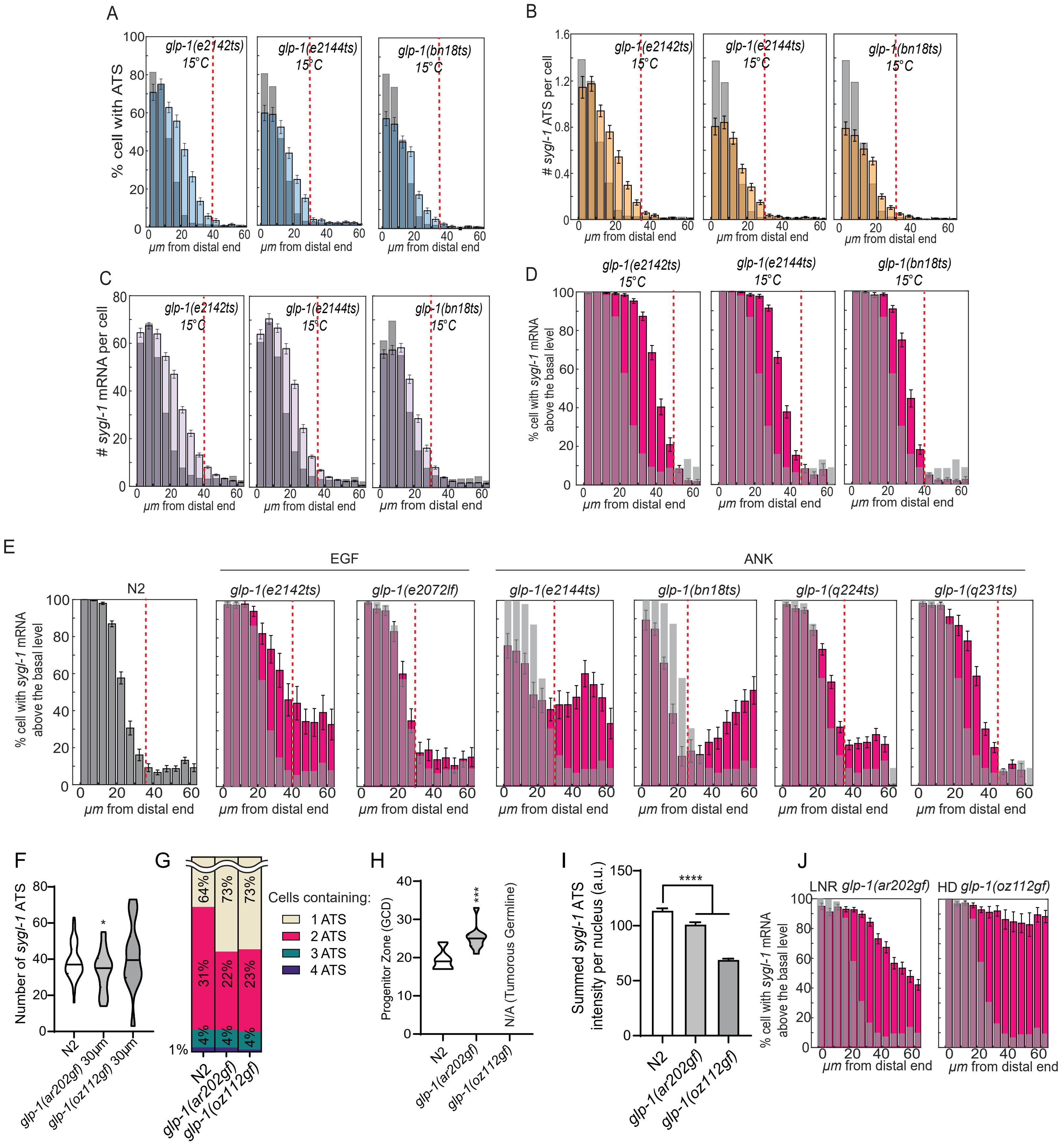
A
% cell with ATS
glp-1(e2142ts) 15°C
glp-1(e2144ts) 15°C
glp-1(bn18ts) 15°C
µm from distal end
B
### sygl-1 ATS per cell
C
### sygl-1 mRNA per cell
D
% cell with sygl-1 mRNA above the basal level
E
EGF
ANK
N2
glp-1(e2142ts)
glp-1(e2072lf)
glp-1(e2144ts)
glp-1(bn18ts)
glp-1(q224ts)
glp-1(q231ts)
F
Number of sygl-1 ATS
N2
glp-1(ar202gf) 30µm
glp-1(oz112gf) 30µm
G
Cells containing:
1 ATS
2 ATS
3 ATS
4 ATS
64%
73%
73%
31%
22%
23%
4%
1%
N2
glp-1(ar202gf)
glp-1(oz112gf)
H
Progenitor Zone (GCD)
N/A (Tumorous Germline)
***
I
Summed sygl-1 ATS intensity per nucleus (a.u.)
****
J
LNR glp-1(ar202gf)
HD glp-1(oz112gf)

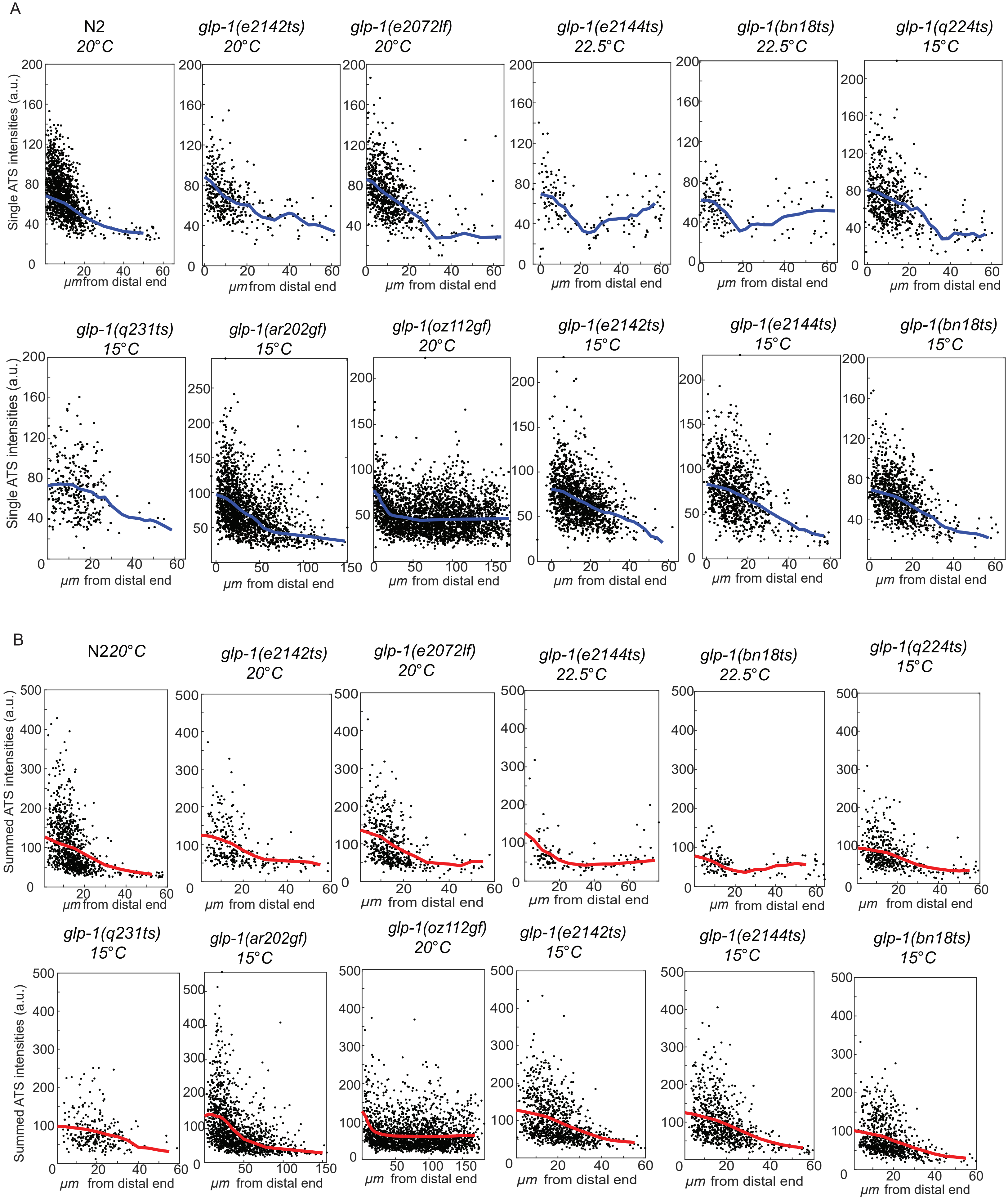
A
N2
20°C
glp-1(e2142ts)
20°C
glp-1(e2072lf)
20°C
glp-1(e2144ts)
22.5°C
glp-1(bn18ts)
22.5°C
glp-1(q224ts)
15°C
glp-1(q231ts)
15°C
glp-1(ar202gf)
15°C
glp-1(oz112gf)
20°C
glp-1(e2142ts)
15°C
glp-1(e2144ts)
15°C
glp-1(bn18ts)
15°C
Single ATS intensities (a.u.)
μm from distal end
B
N220°C
glp-1(e2142ts)
20°C
glp-1(e2072lf)
20°C
glp-1(e2144ts)
22.5°C
glp-1(bn18ts)
22.5°C
glp-1(q224ts)
15°C
glp-1(q231ts)
15°C
glp-1(ar202gf)
15°C
glp-1(oz112gf)
20°C
glp-1(e2142ts)
15°C
glp-1(e2144ts)
15°C
glp-1(bn18ts)
15°C
Summed ATS intensities (a.u.)
μm from distal end

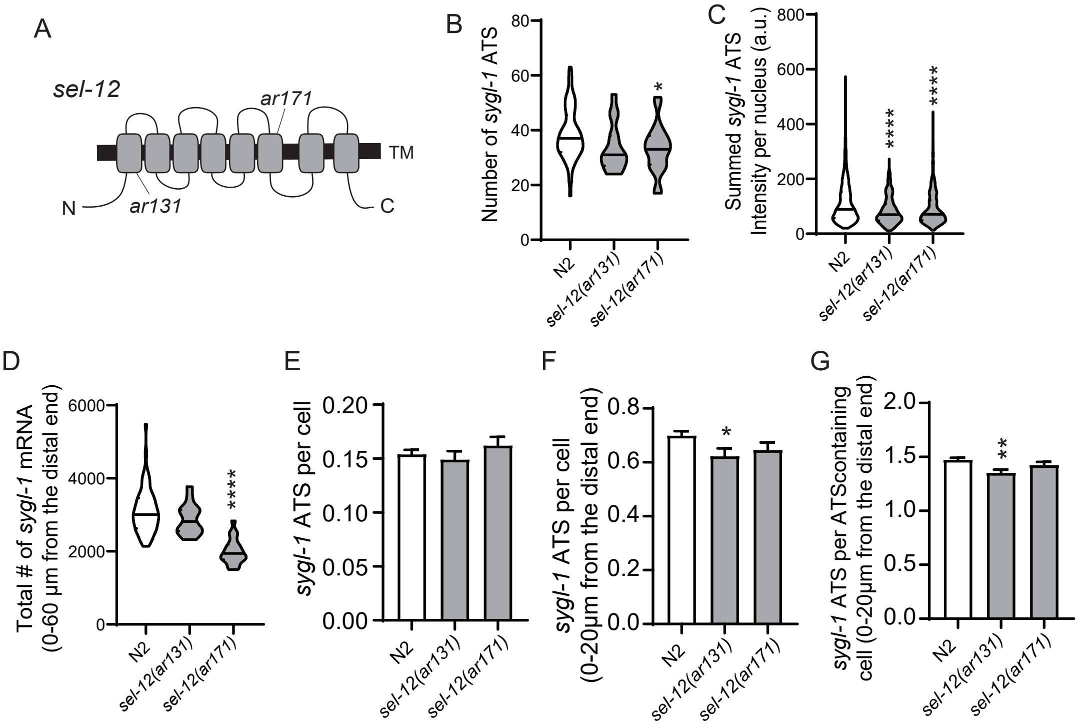

A
sel-12
ar171
ar131
TM
N
C
B
Number of sygl-1 ATS
80
60
40
20
0
*
N2
sel-12(ar131)
sel-12(ar171)
C
Summed sygl-1 ATS Intensity per nucleus (a.u.)
800
600
400
200
0
****
****
N2
sel-12(ar131)
sel-12(ar171)
D
Total # of sygl-1 mRNA (0-60 µm from the distal end)
6000
4000
2000
0
****
N2
sel-12(ar131)
sel-12(ar171)
E
sygl-1 ATS per cell
0.20
0.15
0.10
0.05
0.00
N2
sel-12(ar131)
sel-12(ar171)
F
sygl-1 ATS per cell (0-20µm from the distal end)
0.8
0.6
0.4
0.2
0.0
*
N2
sel-12(ar131)
sel-12(ar171)
G
sygl-1 ATS per ATScontaining cell (0-20µm from the distal end)
2.0
1.5
1.0
0.5
0.0
**
N2
sel-12(ar131)
sel-12(ar171)

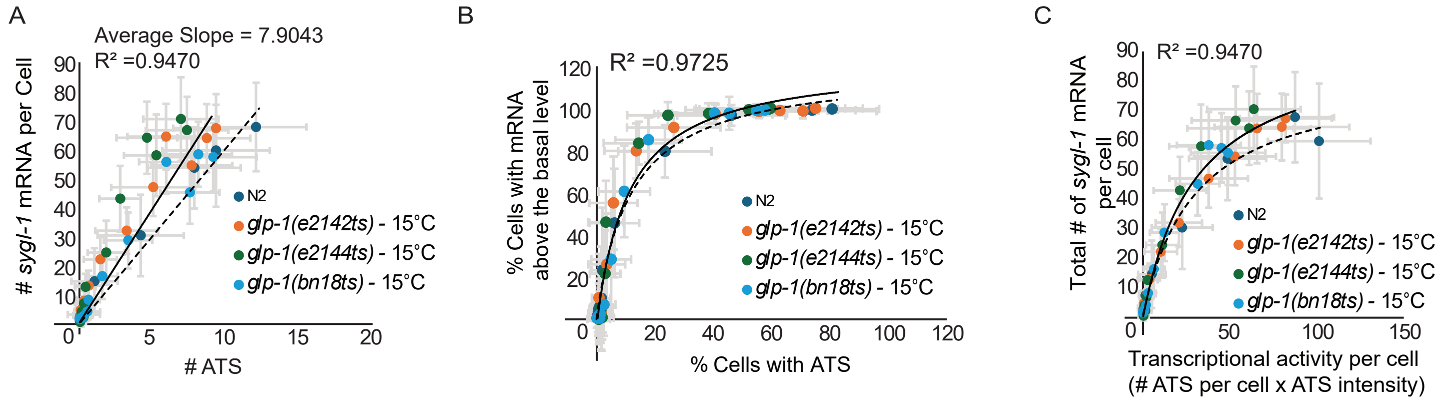

A
Average Slope = 7.9043
R² =0.9470
### sygl-1 mRNA per Cell
### ATS
N2
glp-1(e2142ts) - 15°C
glp-1(e2144ts) - 15°C
glp-1(bn18ts) - 15°C
B
R² =0.9725
% Cells with mRNA above the basal level
% Cells with ATS
N2
glp-1(e2142ts) - 15°C
glp-1(e2144ts) - 15°C
glp-1(bn18ts) - 15°C
C
R² =0.9470
Total # of sygl-1 mRNA per cell
Transcriptional activity per cell
(# ATS per cell x ATS intensity)
N2
glp-1(e2142ts) - 15°C
glp-1(e2144ts) - 15°C
glp-1(bn18ts) - 15°C

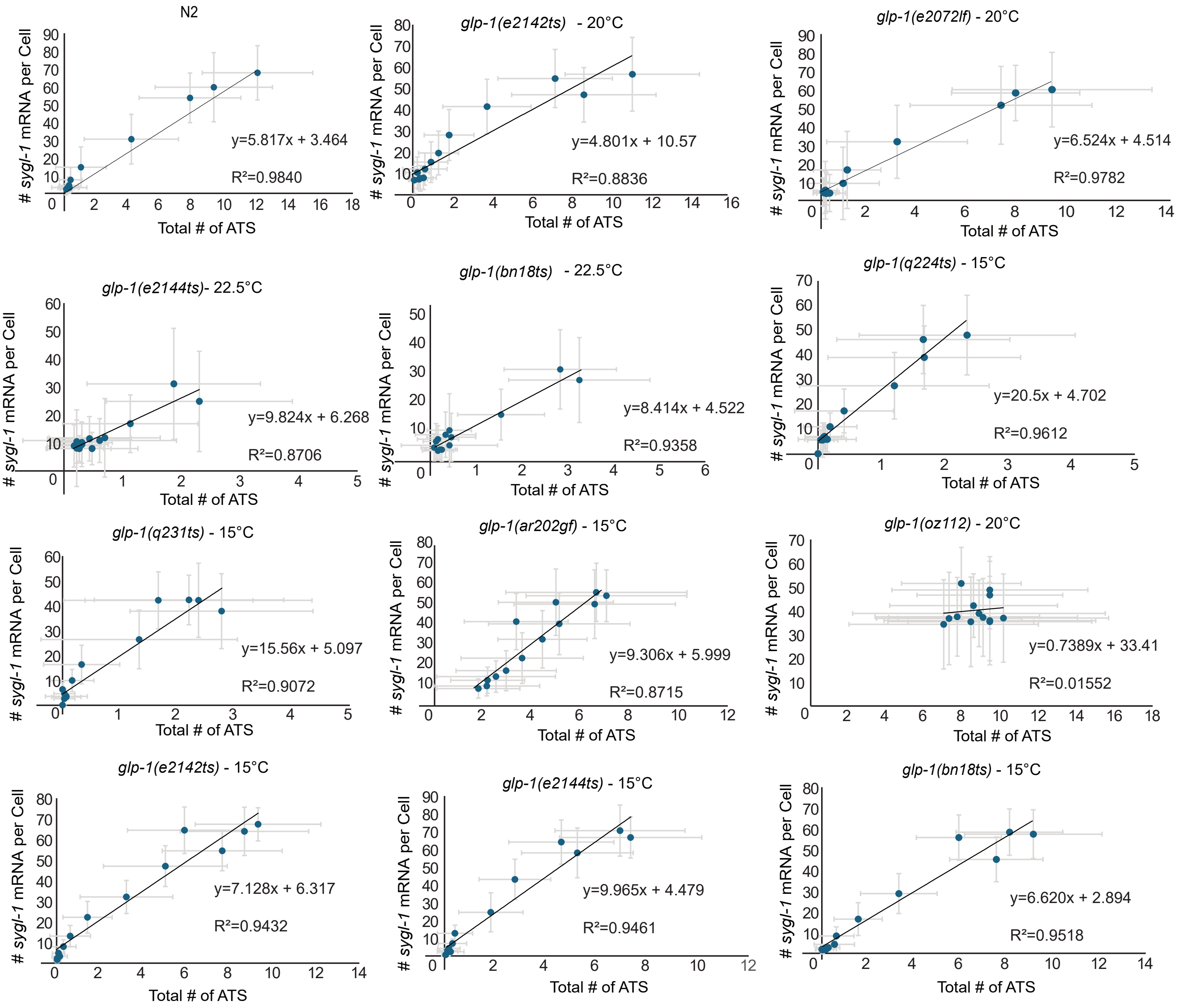

N2
### *sygl-1* mRNA per Cell
y=5.817x + 3.464
R²=0.9840
Total # of ATS
*glp-1(e2142ts)* - 20°C
### *sygl-1* mRNA per Cell
y=4.801x + 10.57
R²=0.8836
Total # of ATS
*glp-1(e2072lf)* - 20°C
### *sygl-1* mRNA per Cell
y=6.524x + 4.514
R²=0.9782
Total # of ATS
*glp-1(e2144ts)*- 22.5°C
### *sygl-1* mRNA per Cell
y=9.824x + 6.268
R²=0.8706
Total # of ATS
*glp-1(bn18ts)* - 22.5°C
### *sygl-1* mRNA per Cell
y=8.414x + 4.522
R²=0.9358
Total # of ATS
*glp-1(q224ts)* - 15°C
### *sygl-1* mRNA per Cell
y=20.5x + 4.702
R²=0.9612
Total # of ATS
*glp-1(q231ts)* - 15°C
### *sygl-1* mRNA per Cell
y=15.56x + 5.097
R²=0.9072
Total # of ATS
*glp-1(ar202gf)* - 15°C
### *sygl-1* mRNA per Cell
y=9.306x + 5.999
R²=0.8715
Total # of ATS
*glp-1(oz112)* - 20°C
### *sygl-1* mRNA per Cell
y=0.7389x + 33.41
R²=0.01552
Total # of ATS
*glp-1(e2142ts)* - 15°C
### *sygl-1* mRNA per Cell
y=7.128x + 6.317
R²=0.9432
Total # of ATS
*glp-1(e2144ts)* - 15°C
### *sygl-1* mRNA per Cell
y=9.965x + 4.479
R²=0.9461
Total # of ATS
*glp-1(bn18ts)* - 15°C
### *sygl-1* mRNA per Cell
y=6.620x + 2.894
R²=0.9518
Total # of ATS

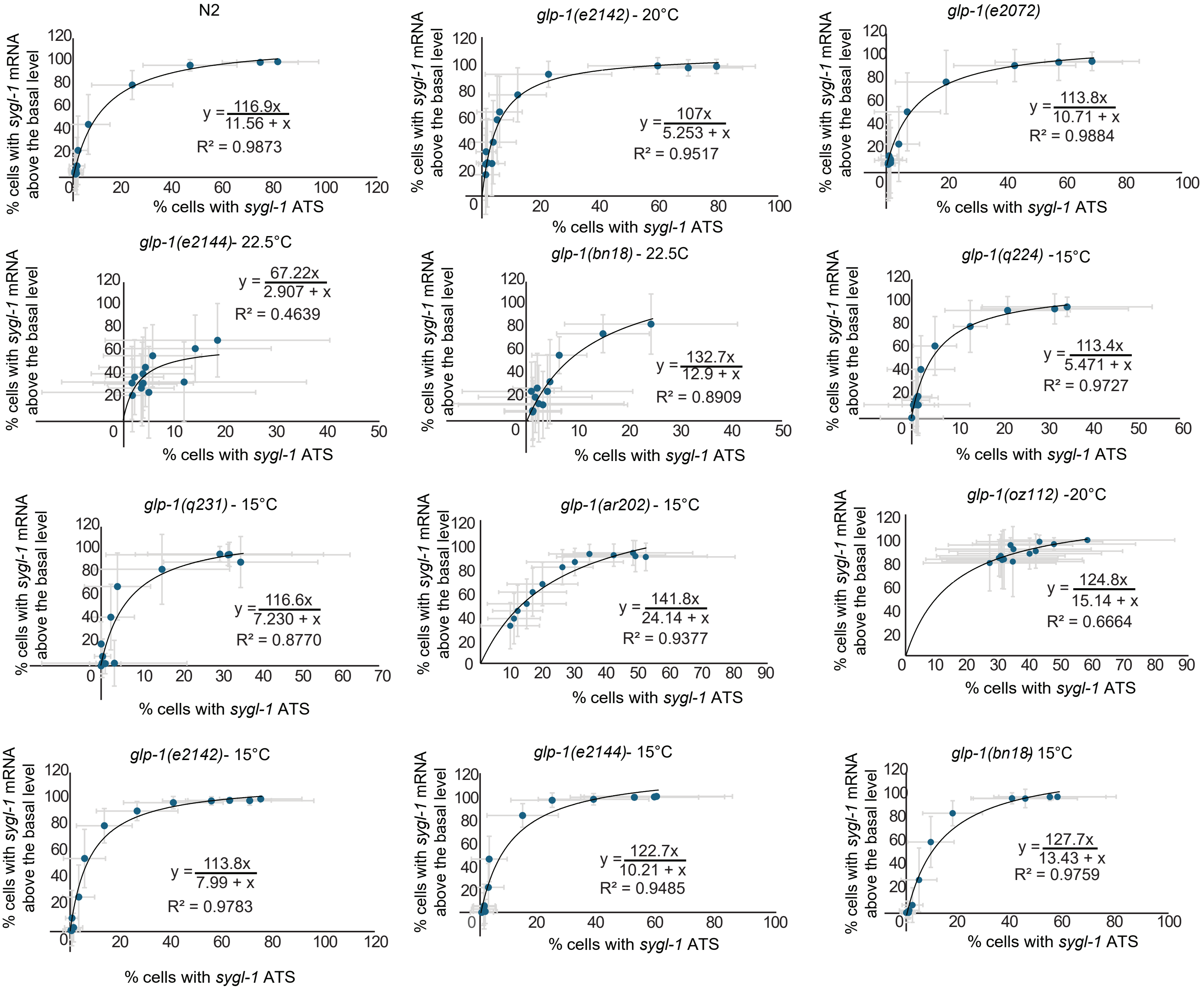

N2
y = 116.9x / 11.56 + x
R² = 0.9873
glp-1(e2142) - 20°C
y = 107x / 5.253 + x
R² = 0.9517
glp-1(e2072)
y = 113.8x / 10.71 + x
R² = 0.9884
glp-1(e2144)- 22.5°C
y = 67.22x / 2.907 + x
R² = 0.4639
glp-1(bn18) - 22.5C
y = 132.7x / 12.9 + x
R² = 0.8909
glp-1(q224) -15°C
y = 113.4x / 5.471 + x
R² = 0.9727
glp-1(q231) - 15°C
y = 116.6x / 7.230 + x
R² = 0.8770
glp-1(ar202) - 15°C
y = 141.8x / 24.14 + x
R² = 0.9377
glp-1(oz112) -20°C
y = 124.8x / 15.14 + x
R² = 0.6664
glp-1(e2142)- 15°C
y = 113.8x / 7.99 + x
R² = 0.9783
glp-1(e2144)- 15°C
y = 122.7x / 10.21 + x
R² = 0.9485
glp-1(bn18) 15°C
y = 127.7x / 13.43 + x
R² = 0.9759
% cells with sygl-1 mRNA above the basal level
% cells with sygl-1 ATS

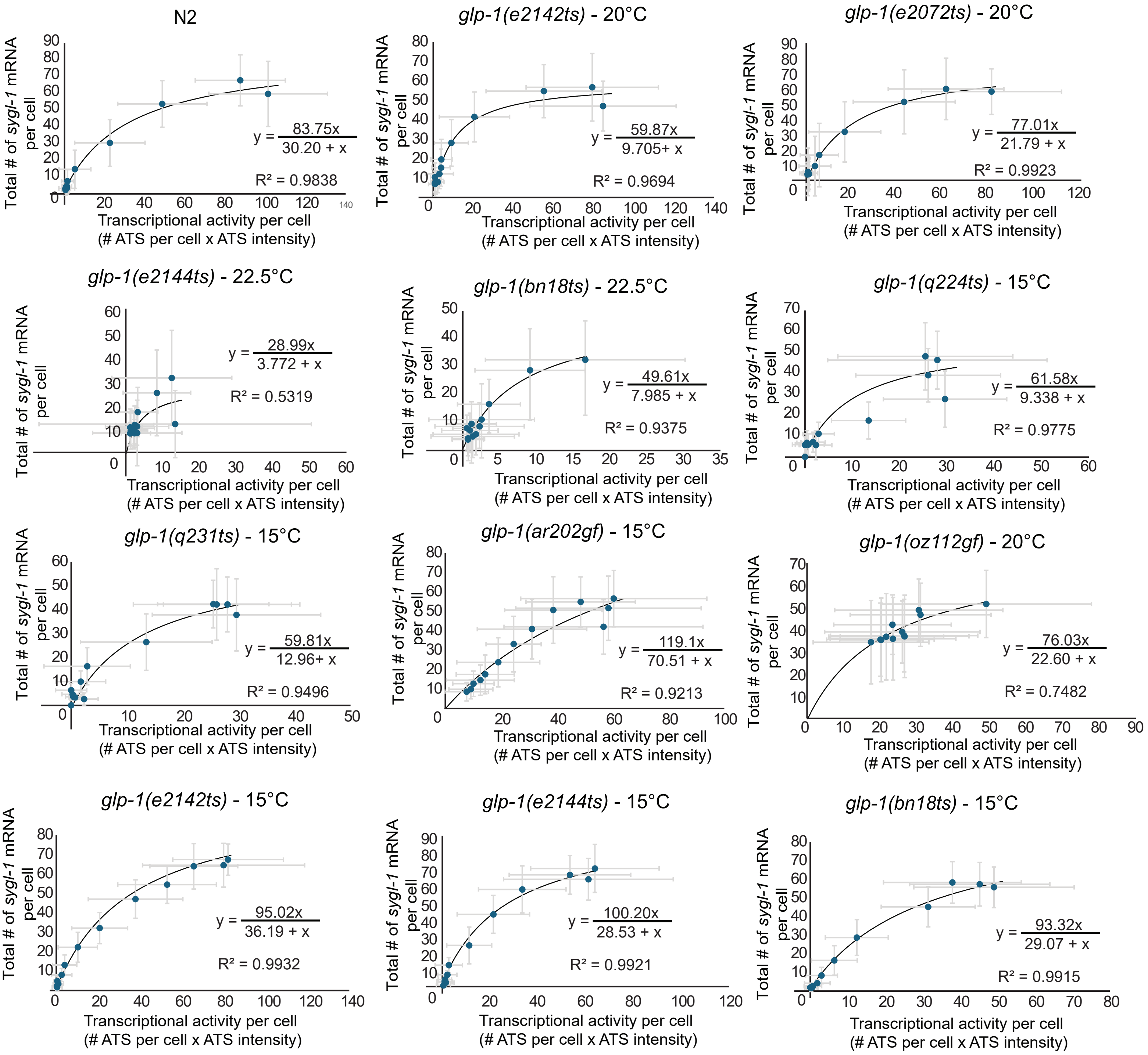
N2
Total # of sygl-1 mRNA per cell
y = 83.75x / 30.20 + x
R² = 0.9838
Transcriptional activity per cell (# ATS per cell x ATS intensity)
glp-1(e2142ts) - 20°C
y = 59.87x / 9.705+ x
R² = 0.9694
glp-1(e2072ts) - 20°C
y = 77.01x / 21.79 + x
R² = 0.9923
glp-1(e2144ts) - 22.5°C
y = 28.99x / 3.772 + x
R² = 0.5319
glp-1(bn18ts) - 22.5°C
y = 49.61x / 7.985 + x
R² = 0.9375
glp-1(q224ts) - 15°C
y = 61.58x / 9.338 + x
R² = 0.9775
glp-1(q231ts) - 15°C
y = 59.81x / 12.96+ x
R² = 0.9496
glp-1(ar202gf) - 15°C
y = 119.1x / 70.51 + x
R² = 0.9213
glp-1(oz112gf) - 20°C
y = 76.03x / 22.60 + x
R² = 0.7482
glp-1(e2142ts) - 15°C
y = 95.02x / 36.19 + x
R² = 0.9932
glp-1(e2144ts) - 15°C
y = 100.20x / 28.53 + x
R² = 0.9921
glp-1(bn18ts) - 15°C
y = 93.32x / 29.07 + x
R² = 0.9915
